# Polyploid cancer cells reveal signatures of chemotherapy resistance

**DOI:** 10.1101/2024.08.19.608632

**Authors:** Michael J. Schmidt, Amin Naghdloo, Rishvanth K. Prabakar, Mohamed Kamal, Radu Cadaneanu, Isla P. Garraway, Michael Lewis, Ana Aparicio, Amado Zurita-Saavedra, Paul Corn, Peter Kuhn, Kenneth J. Pienta, Sarah R. Amend, James Hicks

**Affiliations:** Convergent Science Institute in Cancer, Michelson Center for Convergent Bioscience, University of Southern California, Los Angeles, California, USA; Currently at: Simons Center for Quantitative Biology, Cold Spring Harbor Laboratory, Cold Spring Harbor, NY, USA; Department of Zoology, Faculty of Science, Benha University, Benha, Egypt; Department of Urology, Jonsson Comprehensive Cancer Center, David Geffen School of Medicine at UCLA and VA Greater Los Angeles, University of California, Los Angeles, Los Angeles, California, USA; VA Greater Los Angeles Medical Center, Los Angeles, CA, USA; Departments of Medicine and Pathology, Cedars-Sinai Medical Center, Los Angeles, CA, USA; Center for Cancer Research and Cellular Therapeutics, Clark, Atlanta, GA, USA; Department of Genitourinary Medical Oncology, The University of Texas MD Anderson Cancer Center, Houston, TX, USA; Cancer Ecology Center, The Brady Urological Institute, Johns Hopkins University School of Medicine, Baltimore, MD, USA

**Keywords:** polyploid giant cancer cell, circulating tumor cell, progression-free survival, liquid biopsy, polyaneuploid cancer cell state, chemotherapy resistance, single cell

## Abstract

Therapeutic resistance in cancer significantly contributes to mortality, with many patients eventually experiencing recurrence after initial treatment responses. Recent studies have identified therapy-resistant large polyploid cancer cells in patient tissues, particularly in late-stage prostate cancer, linking them to advanced disease and relapse. Here, we analyzed bone marrow aspirates from 44 advanced prostate cancer patients and found the presence of circulating tumor cells with increased genomic content (CTC-IGC) was significantly associated with poorer progression- free survival. Single cell copy number profiling of CTC-IGC displayed clonal origins with typical CTCs, suggesting complete polyploidization. Induced polyploid cancer cells from PC3 and MDA-MB-231 cell lines treated with docetaxel or cisplatin were examined through single cell DNA sequencing, RNA sequencing, and protein immunofluorescence. Novel RNA and protein markers, including HOMER1, TNFRSF9, and LRP1, were identified as linked to chemotherapy resistance. These markers were also present in a subset of patient CTCs and associated with recurrence in public gene expression data. This study highlights the prognostic significance of large polyploid tumor cells, their role in chemotherapy resistance, and their expression of markers tied to cancer relapse, offering new potential avenues for therapeutic development.

## 1. Introduction

While initial treatment efficacy is observed in most patients with prostate or breast cancer, prostate cancers recur in 24-48% of cases [1], and breast cancers relapse in about 30% of patients [2–3]. In general, late-stage metastatic cancers are more difficult to control, and patients are typically treated with chemotherapy; unfortunately, complete response rates from chemotherapy treatments in patients with late stage disease are low [4–5].

Despite defining numerous detailed intrinsic and extrinsic mechanisms that enable cancer cell survival under therapy, therapy resistance remains responsible for over 90% of cancer related deaths [6–8].

Large polyploid tumor cells are correlated with late disease stages, poor prognosis, and therapy resistance across virtually every tumor type [9–13]. Large polyploid tumor cells are induced through various stressors, including common chemotherapies such as docetaxel and cisplatin [14–17]. Evidence has shown that whole genome doubling (WGD) events and altered ploidy levels are poor prognostic indicators across cancer types and are ultimately thought to provide cancer cells the ability to evolve and survive therapy [18–21].

Recent studies have shown that large polyploid tumor cells can give rise to viable progeny that display more malignant and stem cell characteristics than the parental population they descended from [22]. Importantly, targeting identified pathways, including AP-1, HIF2a, cholesterol-related, and embryogenic- related pathways, reduced the number of surviving large polyploid cancer cells, as well as surviving progeny cells following therapy [22–26]. While significant, these studies lack single-cell molecular resolution and note that not all cells are eliminated. What ultimately matters is that some cancer cells are still capable of survival and result in disease progression. Identification of novel biomarkers that can predict patients’ recurrence and resistance to therapy may lead to better treatment outcomes.

We find that the presence of circulating tumor cells (CTCs) with increased genomic content in the bone marrow aspirate of late-stage prostate cancer patients is significantly associated with worse progression free survival. We comprehensively evaluated large polyploid tumor cells (prostate cancer PC3 and breast cancer MDA-MB-231) that survive following treatment with two chemotherapy classes (cisplatin and docetaxel), and functionally characterize the surviving cells through a multi-omic approach, including morphometric, genomic, and transcriptomic profiling at the single cell level. We find that progeny cells differed substantially from the original parental population and most closely resembled the transcriptome of the large polyploid tumor cells from which they were derived. We also find novel markers associated with chemotherapy survival are upregulated in cells that survive treatment, are retained in the progeny from surviving cells, and are significantly associated with recurrence in prostate and breast cancer at the RNA level. These novel survival biomarkers are expressed at the protein level in the CTCs of patients who also have recurrent disease. Taken together, our results highlight novel biomarkers of survival and shed light on the functionality of large polyploid tumor cells and their role in disease recurrence.

## 2. Methods

### Patient sample collection and processing

Liquid biopsy samples were collected from clinical sites and processed at the University of Southern California as previously described [27–28]. Briefly, peripheral blood (PB) and bone marrow aspirate (BM) samples were collected from patients immediately starting treatment on trial NCT01505868 that evaluated cabazitaxel with or without carboplatin in patients with metastatic castration-resistant prostate cancer. Samples were collected at MD Anderson Cancer Center prior to therapy.

Patients 1 and 3 did not participate in NCT01505868.

Patient 1, a previous case study, was acquired from the Greater Los Angeles Veterans’ Affairs Healthcare System [29]. The bone marrow sample was collected at the time of diagnostic biopsy, prior to treatment. Patient 3, another previous case study [30], was acquired from MD Anderson. All patients gave written informed consent in accordance with approved institutional review board and research development (VA) protocols.

Following isotonic erythrocyte lysis, the entire nucleated fraction was plated onto custom cell adhesion glass slides (Marienfield, Lauda, Germany) and stored at -80°C until use [28].

### Cell culture and drug treatment

PC3 and MDA-MB-231 cell lines were purchased from ATCC and grown in RPMI and DMEM, respectively, with 10% FBS and 0.5% penicillin / streptavidin. Cells were plated at a density of 625,000 cells per T-75 flask. PC3 cells were treated with 5 nM docetaxel (PC3: 5nM, MDA-MB-221: 10nM) or cisplatin (10 µM) for 72 hours. Cells were then allowed to recover in normal medium for 1 or 10 days. When indicated, PC3 cells were re-treated at day 10 post treatment removal. Cells were lifted from culture and plated on Marienfield glass slides for imaging or single cell isolation. All cell line experiments were conducted in triplicate.

To isolate progeny cells, PC3 cells 10 days post treatment were lifted with 1x versene. Biosorter (UnionBio, Holliston, MA) was used to sort single cells based on size and the largest 15% of cells were sorted into ten 96-well plates (n=960 individual wells) containing RPMI medium and then placed in a 37°C incubator.

Media was changed every 2-3 days.

### Immunofluorescent staining

Patient slides in Figure 1 were fixed with paraformaldehyde and stained with a pan-cytokeratin cocktail mixture (see supplementary methods), conjugated mouse anti-human CD45 Alexa Fluor 647 (clone: F10-89-4, MCA87A647, AbD Serotec, Raleigh, NC, USA), Vimentin (Alexa Fluor 488 rabbit IgG monoclonal antibody (Cell Signaling Technology; Cat# 9854BC; Clone: D21H), and 4’,6-diamidino-2-phenylindole (DAPI; D1306, Thermo, Waltham, MA, USA) as previously described [28]. EpCAM (Thermo, 14-9326-82) was included in the pan-cytokeratin cocktail mixture to make an “EPI-cocktail”.

**Figure 1:**
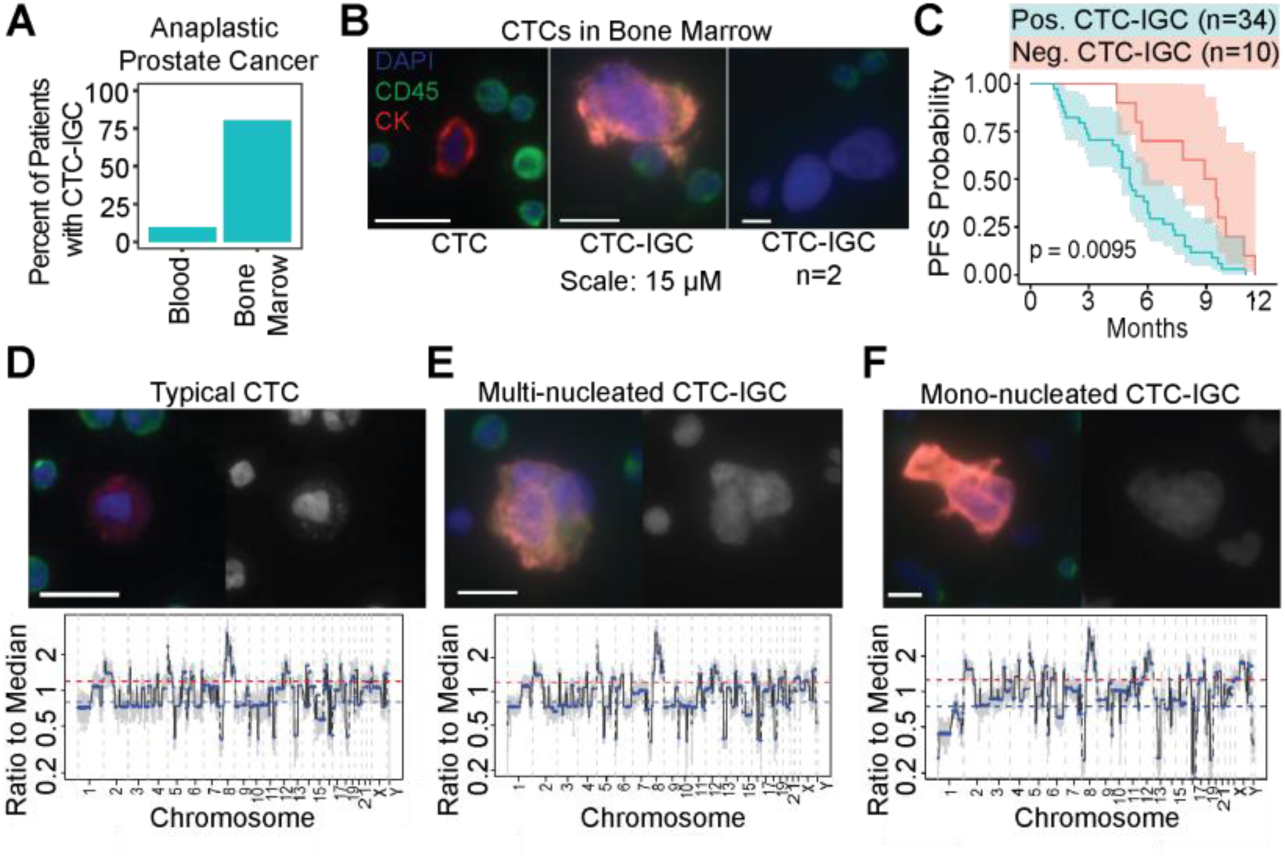
Large tumor cells are found in BM of late-stage prostate cancer patients. (A) Enumeration of patients with matched blood and bone marrow samples with at least 1 CTC-IGC present in liquid biopsy. (B) Representative images of CTC and CTC-IGC found in BM aspirate. Scale bars set to 15 µM. (C) PFS from patients with or without CTC-IGC found in BM samples. (D) Representative image of typical CTC found in BM with merged and DAPI channels (top) and its genomic copy number profile (bottom). (E) Representative image of CTC-IGC found in bone marrow with merged and DAPI images (top) and its genomic copy number profile (bottom). (F) Representative image of mono- nucleated CTC-IGC found in bone marrow with merged and DAPI images (top) and its genomic copy number profile (bottom).

TNFRSF9 (Thermo, PA5-98296) and HOMER1 (Thermo, PA5-21487) primary antibodies were incubated on slides overnight at 4°C with the EPI-cocktail of antibodies. Slides were then washed and incubated at room temperature for two hours with Alexa Fluor 555 goat anti-mouse IgG1 antibody (Thermo, A21127), Alexa Fluor 488 goat anti-rabbit (Thermo, A11034), CD45, and DAPI.

LRP1 (Thermo, 377600) was generated in mice and was therefore not compatible with the EPI-cocktail. Instead, LRP1 was incubated overnight at 4°C. Slides were then washed and incubated at room temperature for two hours with Alexa Fluor 555 goat anti-mouse IgG1 antibody. Next, pre-conjugated Alexa Fluor 488 pan-cytokeratin (53-9003-82, Thermo) recognizing CK 10, 14, 15, 16, and 19 was incubated with conjugated mouse anti-human CD45, and DAPI.

### Slide imaging and analysis

Slides were imaged with an automated high throughput microscope equipped with a 10x optical lens, as previously described [27]. Immunofluorescent and bright field images were collected. Image analysis tool, available at https://github.com/aminnaghdloo/slide-image-utils, was developed in python using the OpenCV and scikit-image packages [31–32]. Briefly, each fluorescent channel was segmented individually using adaptive thresholding and merged into one cell mask. Cell mask and DAPI mask were used to extract features and fluorescent intensity statistics of single cells and their nuclei, respectively. For nucleus size analysis, equivalent diameter was calculated from nucleus area, assuming a circular shape.

### Fluorescence *in situ* Hybridization

Probes for centromeres of chromosomes 1 (CHR01-10-GR) and 10 (CHR10-10-GR) were purchased from Empire Genomics (Depew, New York) and the hybridization was carried out on Marienfeld glass slides per the manufacturer’s instructions. Slides were then stained with DAPI and then imaged.

### Single cell copy number profiling

Single cells were isolated as previously described [28]. Copy number profiling from low pass whole genome sequencing samples was conducted as previously described (see supplementary methods) [33–34].

### Single cell RNA sequencing

Single cells were isolated and picked via micro-manipulation as previously described. RNA was extracted via a modified Smart-Seq2 approach and library prepped with Nextera XT (Illumina, San Diego, CA). Cells were sequenced paired end by 150 base-pairs on an Illumina HiSeq 4000 (Fulgent). Read adapters were trimmed with TrimGalore (version 0.6.7) and aligned with the HiSat2 (v2.2.1). Picard (v3.0.0) was used to visualize RNA mapping quality control [35–37]. HTSeq (v2.0.2) was used to generate a gene count matrix [38].

The SingleCellExperiment package (v4.2.2) was utilized for inputting count data into downstream analyses, such as converting to Seurat (v4.3.0) and edgeR (v3.36.0) count matrices [39]. Downstream analysis was performed with R (v4.1.2). Data visualization was performed with Seurat and ggplot2 (v3.4.4), and Pheatmap (v1.0.12) packages.

The edgeRQLFDetRate differential expression pipeline was used to find common upregulated genes in polyploid cancer cells [40]. Sequencing batches were controlled for. Shared genes expressed in surviving large cells were intersected through R.

Gene datasets were downloaded directly from CHEA3 [41] and MSigDB [42] for transcription factor and hallmark pathway enrichment, respectively. Single cell enrichment was conducted through JASMINE [43].

### Survival analysis

Survival analysis from patient bone marrow and peripheral blood samples was performed with the Survival R package (v3.5.5) and plotted with ggplot2 (v3.4.4). Public gene expression survival analysis was analyzed via PanCancSurvPlot [44] for prostate cancer (GSE116918) and breast cancer (GSE10893) [45–46].

## 3. Results

### CTCs with increased genomic content (CTC-IGC) are found in the bone marrow of late-stage prostate cancer patients and are correlated with worse progression free survival

Liquid biopsies from peripheral blood and bone marrow aspirate were acquired from a late-stage prostate cancer cohort (NCT01505868). Matched bone marrow and peripheral blood samples from 31 patients were analyzed for CTCs. CTCs with increased genomic content (CTC-IGC), identified as having a nuclear diameter at least double the average of the CTC cell population, were found in 9.7% of peripheral blood samples. CTC- IGC were present in 80.6% of bone marrow samples from the same patients (Fig. 1A-B, S1). Survival analysis with 44 bone marrow samples (from the 31 patients with matched blood samples and 13 patients without matched blood) showed that presence of at least one CTC-IGC detected in the bone marrow was associated with decreased progression-free survival (Fig. 1C, Table S1). Previous treatment history was available for 33 of the 44 patients and primarily included anti-androgens and other hormonal treatments (i.e., bicalutamide, nilutamide, enzalutamide). The six patients who were previously treated with docetaxel were all positive for CTCs-IGC in the bone marrow.

Clonal tumor lineage measured via copy number ratio analysis was confirmed in both typical CTCs and CTCs-IGC. No apparent differences in copy number ratios were identified between the two CTC groups (Fig. 1D-F, S2, S3). These observations show that CTC-IGC can be found in blood and bone marrow aspirate, are tumor derived, and thus may contribute towards relapse in late-stage prostate cancer. Despite the apparent WGD of CTC-IGC, these cells retain the original tumor copy number profile. To understand the importance and behavior of this phenotype, we used an *in vitro* model of polyploid tumor cells to investigate their relationship with therapeutic resistance.

### Large polyploid cancer cells form as a response to chemotherapy in prostate and breast cancer models

PC3 and MDA-MB-231 cells were treated with sublethal doses of docetaxel or cisplatin for 72 hours. Following chemotherapy, cells were allowed to recover for 1 or 10 days in their regular growth medium, lifted from culture, plated on specialized glass slides, stained with cell and nuclear markers, then imaged through high content scanning and evaluated for nuclear size and other morphometric comparisons (Fig. 2A). While there was significant cell death as expected (Fig. S4A), surviving cells increased in both nuclear diameter and cell size as a function of time (Fig. 2B-E; Fig. S4B).

**Figure 2:**
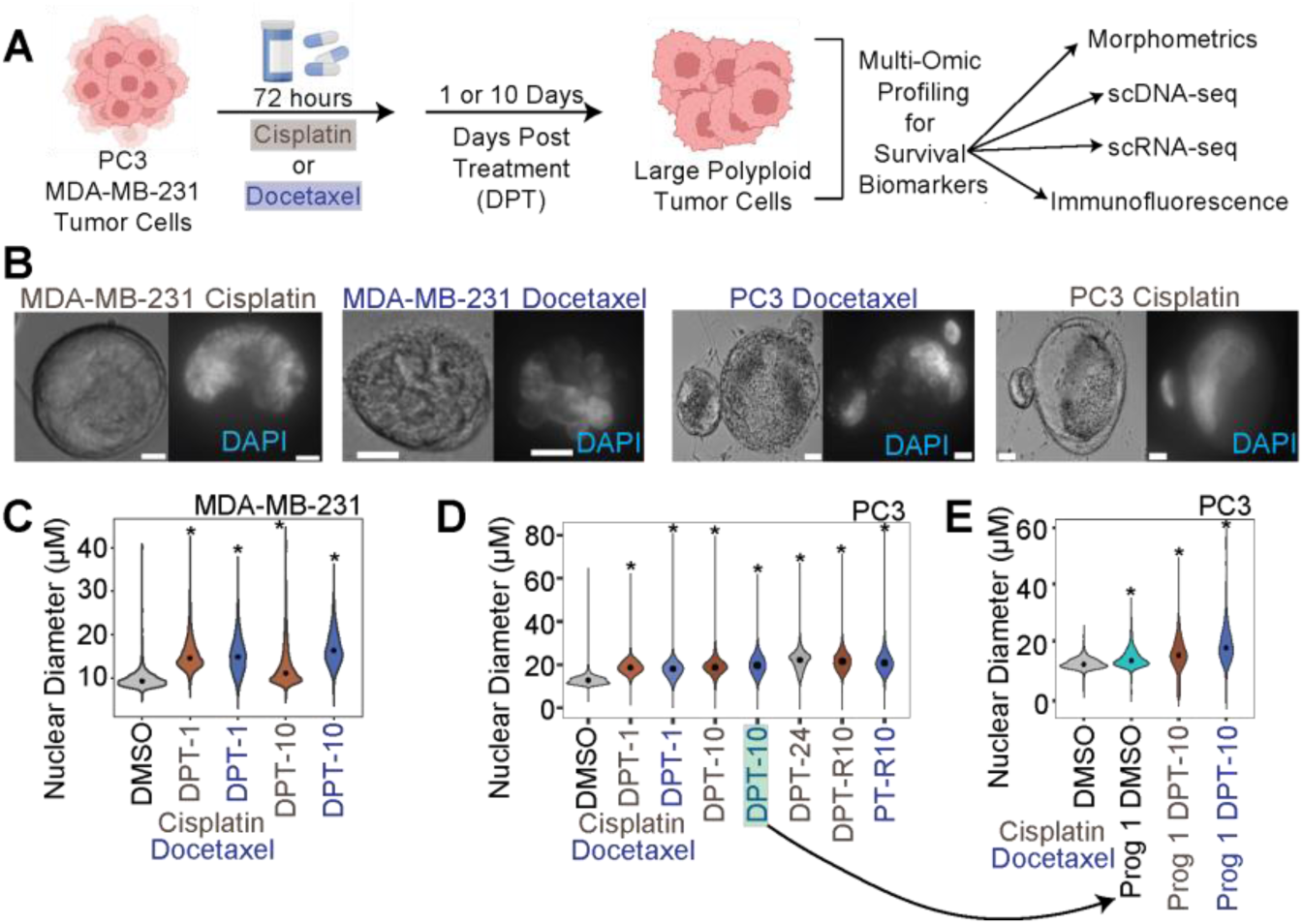
Large polyploid tumor cells are induced following chemotherapy exposure in MDA-MB-231 and PC3 cell lineages. (A) Experimental schematic for in vitro investigation of surviving polyploid cells. (B) Representative bright field (left) and DAPI (right) 40x images of MDA-MB-231 cells and PC3 cells treated with docetaxel and cisplatin (right) 10 days post treatment recovery. (C) Nuclear diameter for all treated conditions for MDA-MB-231 cells. (D) Nuclear diameter for all treated conditions for PC3 cells. Arrow indicates the condition where single cell progeny originated from. (E) Nuclear diameter of PC3 control parental and progeny-1 cells, and progeny-1 10 days post-treatment cells.

To evaluate resistance, we treated cells that survived cisplatin treatment (10 days post treatment; 10 DPT) with cisplatin or docetaxel. Compared to the control condition (initially cisplatin treated and then re-treated with DMSO) cell counts and cell viability were not significantly impacted, suggesting that these cells are not sensitive to additional rounds of chemotherapy (Fig. S4A, S4D, 2E).

To obtain progeny cells from a single chemotherapy- induced surviving polyploid cell, we isolated and single-cell seeded PC3 cells 10 days post-cisplatin release (n=480) and 10 days post-docetaxel release (n=960) and monitored for colony formation. From these, only 2 polyploid docetaxel-treated PC3 cells gave rise to progeny after 2 months (progeny-1) and 2.5 months (progeny-2). Progeny-2 failed to proliferate following the first passage. Over the course of the three-month experiment, approximately 50% of the polyploid cells treated with either cisplatin or docetaxel remained viable and adherent. The dividing progeny-1 cells displayed a larger nuclear and cellular diameter than the parental PC3 population from which it originated (Fig. 2F, S4). We treated progeny-1 with docetaxel or cisplatin and found that the population was sensitive to both chemotherapies. Further, following 10 days of recovery, surviving progeny-1 cells had increased nuclear and cell diameter, similar to what was observed from the original parent population (Fig. 2F, S4B).

### Surviving PC3 polyploid cancer cells show no additional copy number ratio alterations compared to parental controls

To evaluate the presence of genomic alterations in the surviving polyploid cells and their progeny, we assayed copy number status and cell ploidy. Strikingly, surviving large polyploid PC3 and MDA-MB-231 cells from both docetaxel and cisplatin treatments showed no apparent copy number ratio differences compared to control cells (Fig. 3A-C, S5). This result confirms patient data in that copy number ratio status does not differ between CTCs with normal nuclei and CTCs with larger nuclei (Fig. 1D-F) and suggests that cells are undergoing complete WGD rather than displaying specific copy number breakpoints. While copy number status did display minor differences in the progeny-1 compared to parental control (e.g., an increased 3p gain) (Fig. 3A- B), no substantial alterations were observed. Conversely, progeny- 2, the clone that did not survive the first passage, displayed the most aberrant copy number profile compared to the other conditions (i.e., 6 gain and 4p gain) and clustered separately from the other PC3 cell conditions (Fig. 3A).

**Figure 3:**
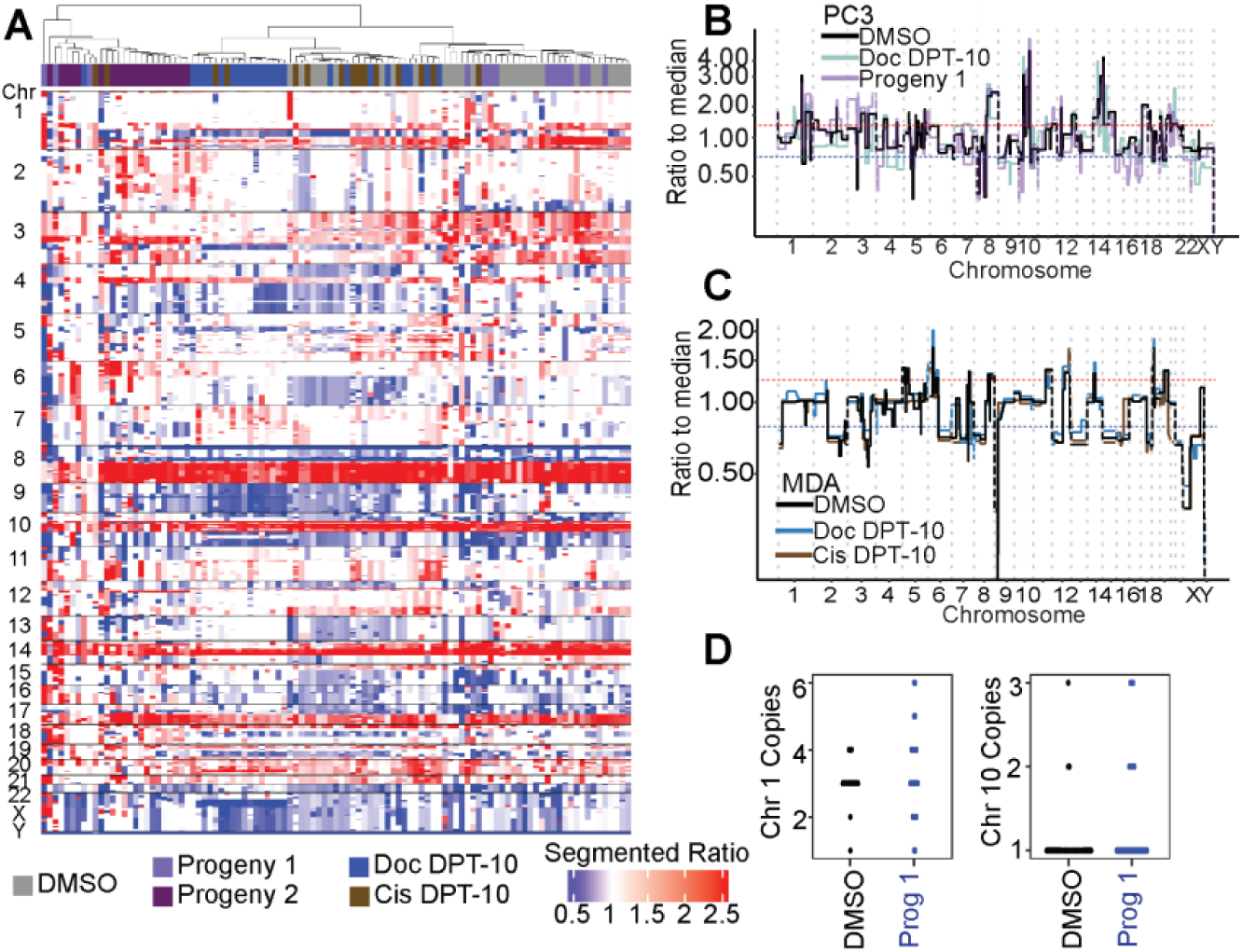
Genomics of PC3 DMSO control, docetaxel treated cells, and docetaxel large cell progeny-1. (A) Segmented copy number ratios for PC3 conditions. Ratio of 1 (white) indicates copy number neutral respective to the entire genome. (B) Representative PC3 DMSO, Doc D10, and Progeny-1 copy number ratio profiles. (C) Representative MDA-MB-231 DMSO control, Doc D10, and Cis D10 copy number ratio profiles. (D) FISH for PC3 DMSO control and progeny-1 cells for centromere of chromosome 1 (ploidy = 3) and chromosome 10 (ploidy = 1).

FISH probes for the centromeres of PC3 chromosome 1 (ploidy = 3) and chromosome 10 (ploidy = 1) showed no statistically significant differences when comparing DMSO parental control cells to progeny-1 cells (Fig. 3D), suggesting that any apparent scars of ploidy reduction were not present. These results prompted investigation into the phenotype of these surviving cells.

### Single cell transcriptomic profiling reveals common genes and pathways upregulated in PC3 and MDA-MB-231 polyploid cells

497 PC3 cells were isolated and sequenced in 5 separate batches (Fig. S6) and included: DMSO control (n=129), 1-day post-cisplatin release (n=78), 10 days post-cisplatin release (n=68), 1-day post-docetaxel release (n=45), 10 days post- docetaxel release (n=118), docetaxel progeny-1 (n=12), docetaxel progeny-2 (n=13). Two batches of 203 MDA-MB-231 cells included: DMSO control (n=43), 1-day post-cisplatin release (n=22), 10 days post-cisplatin release (n=62), 1-day post- docetaxel release (n=24), 10 days post-docetaxel release (n=62) (Fig. S7).

Regardless of treatment, a general spatial separation that was dependent on recovery duration was observed in PC3 and MDA-MB-231 cells (Fig. 4A, S8). To identify convergent phenotypes regardless of tumor type or therapy, we evaluated genes that were upregulated in both PC3 and MDA-MB-231 following either cisplatin or docetaxel treatment. MDA-MB-231 cells 10 days post cisplatin or docetaxel release upregulated 1591 shared genes compared to DMSO control; PC3 cells 10 days post cisplatin or docetaxel treatment upregulated 1178 shared genes compared to DMSO control (LFC > 1.5, FDR < 0.01; Fig. 4B).

**Figure 4:**
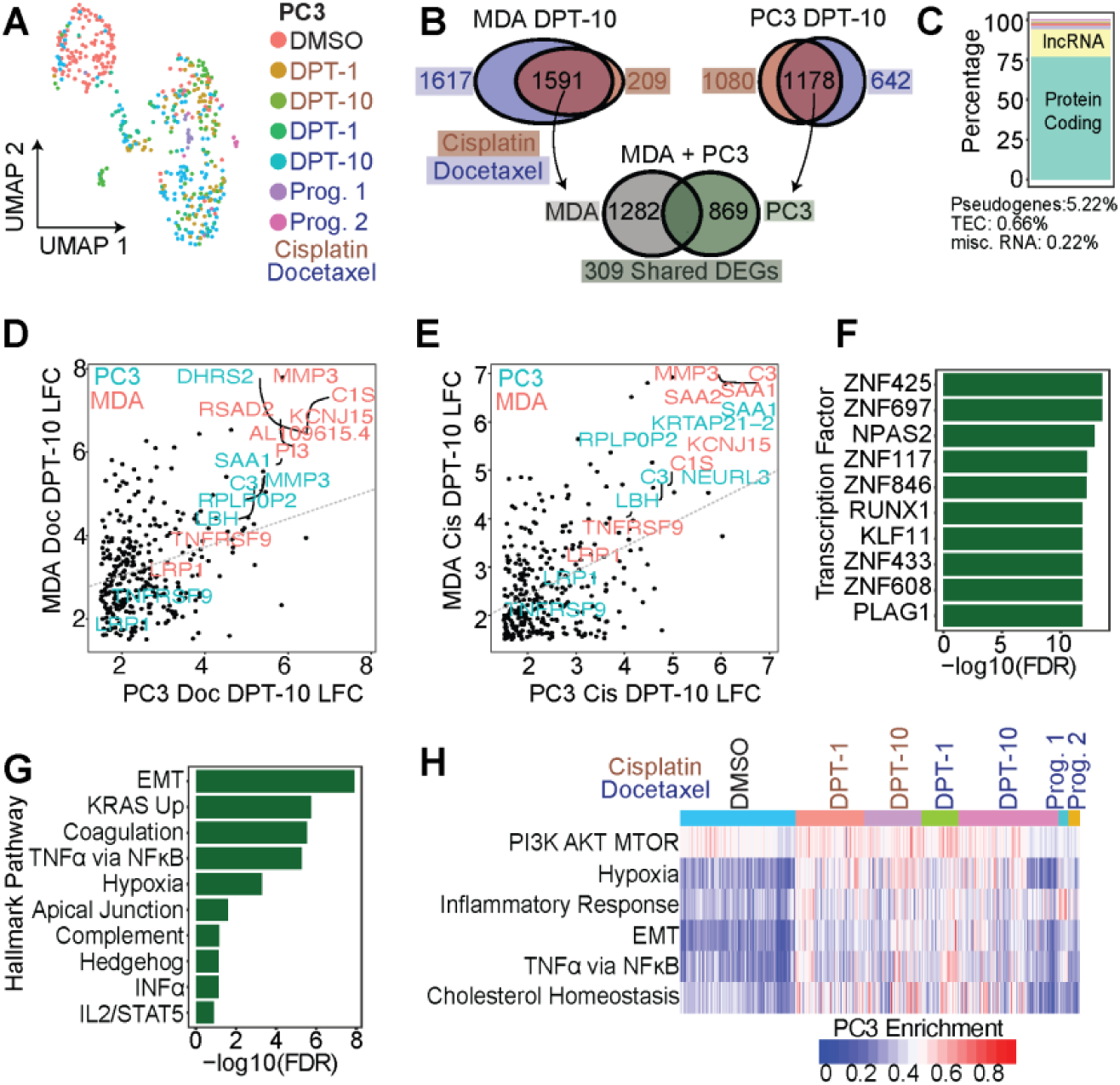
Chemotherapy induced surviving tumor cells share common phenotypes and pathways for survival. (A) UMAP of all conditions for PC3 cells. (B) Comparison of DEGs between MDA- MB-231 large-D10 and PC3 large-D10 cells. Between cisplatin and docetaxel treatments, MDA-MB-231 and PC3 share 309 upregulated genes compared to their respective controls. (B) Genecode annotations for 309 shared genes. (D-E) LFC of shared 309 genes for PC3 vs MDA-MB-231 for docetaxel and cisplatin treatments, respectively. (F) CHEA3 transcription factor enrichment of the shared 309 genes between MDA-MB-231 and PC3 cells. (G) Hallmark enrichment analysis of 309 shared genes. (H) Single cell Hallmark gene set enrichment analysis for all PC3 cells.

Intersection of the shared gene sets showed MDA-MB-231 and PC3 cells that survive either cisplatin or docetaxel exposure shared 309 upregulated genes (Fig. 4B; Table S2). The 309 shared genes were considered a survivor cell enrichment data set, which was further evaluated.

Of the 309 shared genes, 77% were protein coding and 17% were lncRNAs, while the remaining ∼6% were pseudogenes or yet to be experimentally confirmed (TEC, not yet tested; Fig. 4C). Log-fold change values were plotted for PC3-Doc-DPT10 vs MDA-Doc-DPT10 (Fig. 4D) and PC3-Cis-DPT10 vs MDA-Cis-DPT10 (Fig. 4E). Within each treatment class, shared differentially expressed genes (DEGs) were positively correlated between MDA-MB-231 and PC3 cells, indicating the DEGs are upregulated to a similar magnitude.

Common transcription factors (TFs) and hallmark pathways upregulated in the survivors were delineated (Fig. 4F- G). Two significantly enriched TFs, ZNF697 and NPAS2, were previously reported in cells that transition out of senescence and into a proliferative state [47]. Top enriched hallmark pathways in the 309 gene survivor data set were: epithelial-to-mesenchymal transition (EMT), upregulation of KRAS signaling, coagulation, TNFa signaling via NFkB, and hypoxia (Fig. 4F). Single cell gene enrichment confirmed the top upregulated hallmark pathways in the shared survivor data set (Fig. 4H, S8-9). Additional pathways identified to be significantly upregulated in the surviving cells were: PI3K-AKT-mTOR Signaling, Inflammatory Response, and Cholesterol Homeostasis (Fig. 4H, S8-9).

### Identification of HOMER1, TNFRSF9, and LRP1 as putative chemotherapy RNA survival markers

Utilizing the shared cell survivor gene set data, markers were independently evaluated to understand their putative role in chemotherapy survival and polyploid state. All 309 genes were investigated via literature review and queried for terms in September 2023, including: large tumor cell, polyploid giant cancer cell, poly-aneuploid cancer cell, survival pathways, drug resistance, chemotherapy, and apoptosis. With prior knowledge that top upregulated genes (MMP-3, SAA1, and C3) functioned in the execution of apoptosis and clearance of apoptotic bodies (Fig. 4D-E), and that SAA1 and C3 were correlated with better PFS (Fig. S10), they were not considered novel survival markers. The 309 gene survivor cell enrichment data set was also intersected with genes in the top enriched pathways that modulate survival: TNFa via NFkB, PI3K-AKT, and mTOR signaling (Fig. 4G-H). We identified TNFRSF9 and LRP1 as survival biomarkers; these are known to function as cell surface receptors that enhance PI3K activity. This activity, in turn, stimulates AKT, thereby promoting cell survival (Fig. 4D-E, Fig. 5, S11A) [48–50]. Further, we identified HOMER1 as a PC3-specific survival marker (Fig. 5); HOMER1 plays a role in mTOR signaling and protection against apoptosis [51–54].

**Figure 5:**
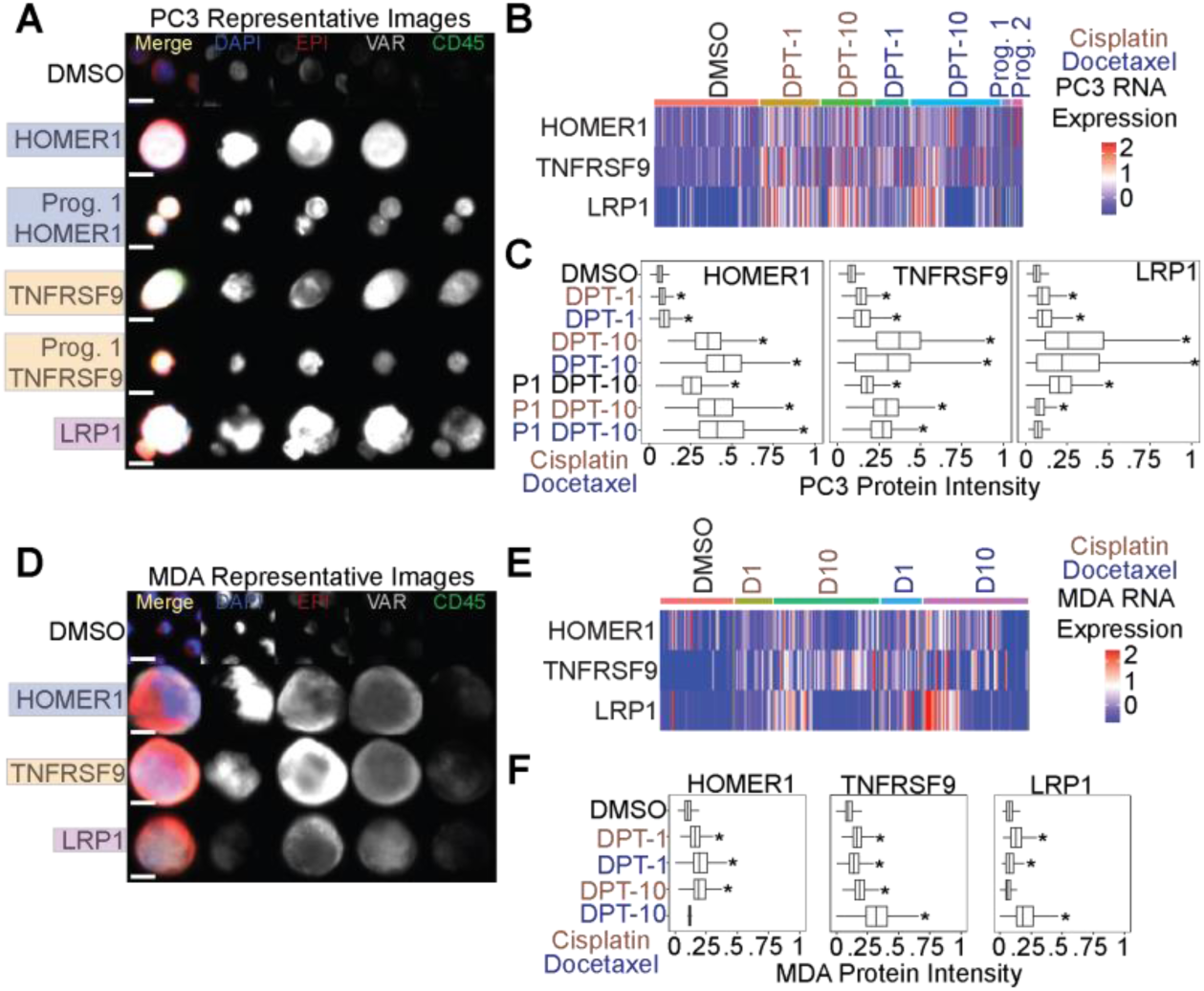
HOMER1, TNFRSF9, and LRP1 are putative markers of chemotherapy resistance. (A) Representative PC3 images of putative marker genes stain in the VAR (4th) channel. DMSO control cells were stained with HOMER1 and were negative. (B) RNA expression for each marker for all PC3 cells. (C) Immunofluorescence quantification for PC3 cells stained with tested markers. (D) Representative MDA-MB-231 images of putative marker genes stained in the VAR (4th) channel. DMSO control cells were stained with HOMER1 and were negative. (E) RNA expression for each marker for all MDA-MB-231 cells. (F) Immunofluorescence quantification for MDA-MB-231 cells stained with tested markers.

### HOMER1, TNFRSF9, and LRP1 are protein markers of chemotherapy survival and are retained in docetaxel treated PC3 progeny

At the protein level, we found surviving PC3 and MDA-MB- 231 cells post-chemotherapy treatment stained positive for HOMER1, TNFRSF9, and LRP1 (Fig. 5A, 5D). Image quantification revealed all PC3 conditions (except for progeny-1 cisplatin day 10 post-treatment release) were significantly upregulated compared to controls (Fig 5A, 5C). Day 10 survivors showed the highest protein expression levels for each marker tested. Importantly, untreated PC3 progeny-1 displayed significantly higher expression in all three survival markers tested compared to parental DMSO control cells, suggesting these markers were retained following treatment (Fig. 5A, 5C). CD45 is typically utilized as a tumor cell exclusion marker that stains for white blood cells. At day 10 post-treatment release time points we noted a gain in CD45 protein expression that was retained in progeny cells in PC3 cells (Fig. 5A, S10). MDA-MB-231 cells also showed a significant upregulation of expression for most markers tested, except HOMER1 for docetaxel day 10 post-treatment release and LRP1 for cisplatin day 10 post-treatment release (Fig. 5D, 5F).

### HOMER1, TNFRSF9, and LRP1 are found at the protein level patient BM samples, and their increased expression is correlated with recurrence in public datasets

A subset of bone marrow samples that displayed a high frequency of CTC-IGC from the prostate cancer patient cohort (Fig. 1) were stained with HOMER1, TNFRSF9, and LRP1 (Fig. 6A). Patient 1 and patient 3 did not participate in the clinical trial but also displayed a high frequency of CTC-IGC (Table 1; see methods). All patients profiled had CTCs that were positive for the tested markers (Fig. 6B). While there were CTC-IGC positive for the marker genes in each patient sample (Fig. 6A), the tested markers were not selective for CTC-IGC (Fig. S12). Patient-5, who displayed the highest percentage of CTCs positive for markers HOMER1 and TNFRSF9, had the shortest PFS at 1.4 months (Fig. 6B, Table 1). Additionally, these survival markers identified cells in the bone marrow that displayed increased genomic content but were negative for canonical epithelial markers (Fig. S13).

**Figure 6:**
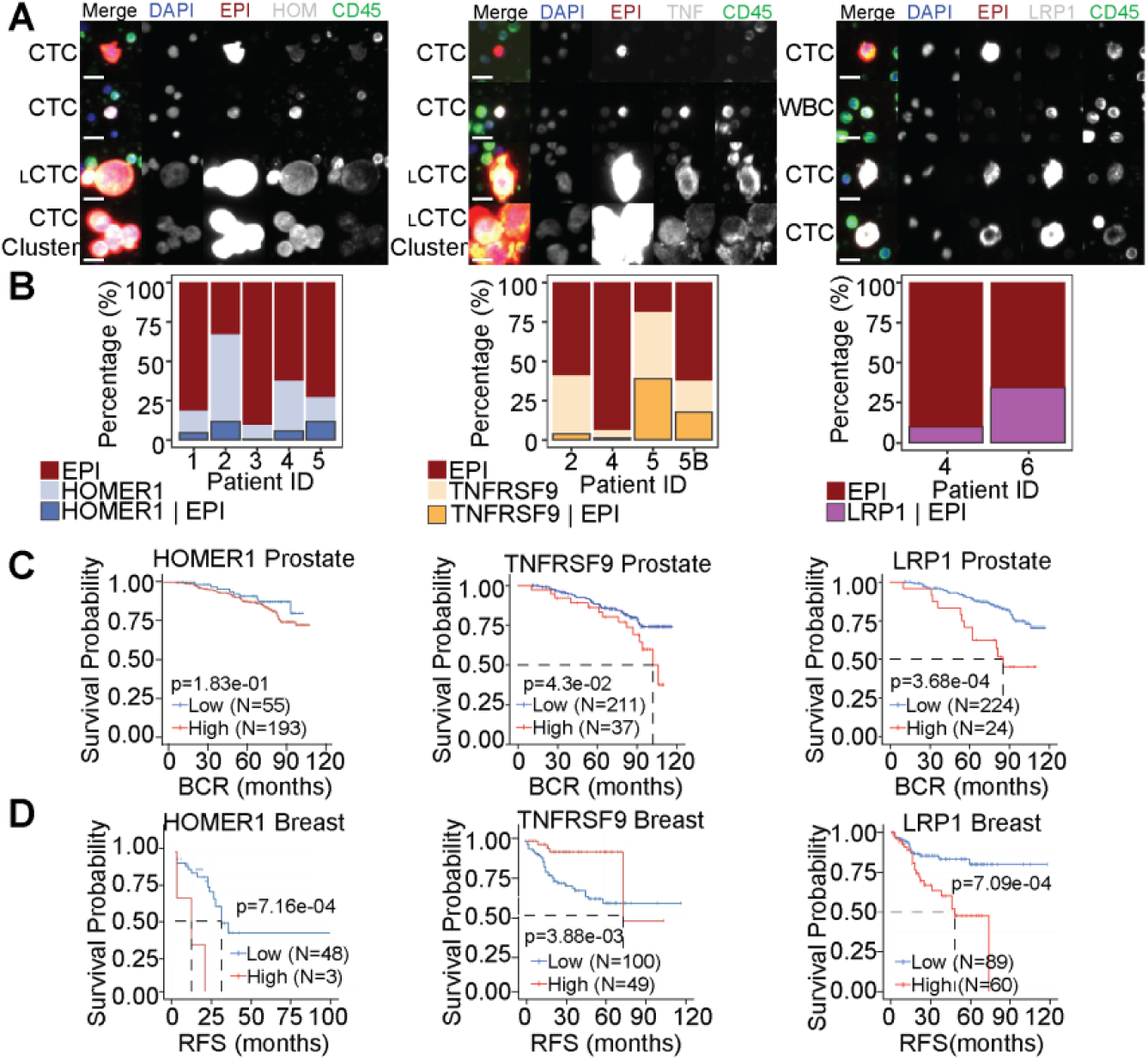
HOMER1, TNFRSF9, and LRP1 are positive on CTCs in the BM aspirate of late-stage prostate cancer and are correlated with recurrence in prostate and breast cancers. (A) Representative CTCs from BM of advanced prostate cancer patients that were stained with survival markers HOMER1 (left), TNFRSF9 (TNF; middle), and LRP1 (right). Tested markers appear as white in the merged image. Scale bars are 15 µM. (B) Percentages of CTCs with EPI positivity and cells that were stained with survival markers HOMER1 (left), TNFRSF9 (middle), and LRP1 (right). Cells that are positive for the marker alone (middle bar) cannot be conclusively labeled a tumor derived cell. LRP1 (right) is also a marker of T-cells, so only cells that were EPI positive were included. (C-D) Kaplan-Meyer survival plots for RNA expression of tested markers in prostate (C) and (D) breast cancer patients.

In publicly available data for previously treated patients, high expression of TNFRSF9 and LRP1 significantly correlated with a shorter progression free survival in patients with prostate cancer; HOMER1 was not statistically significant (p-value = 0.183) (Fig. 6C). High gene expression of TNFRSF9, HOMER1, and LRP1 were all significantly correlated with worse relapse free survival in breast cancer (Fig. 6D). Taken together, we can conclude the survival genes are associated with recurrence at the RNA level and are present on CTCs-IGC in the bone marrow aspirate of late-stage prostate cancer patients.

## 4. Discussion

Our analysis of bone marrow liquid biopsy samples from previously treated advanced prostate cancer patients reveals that the presence of polyploid cancer cells correlates with poorer progression-free survival. Although clinical reports have frequently observed polyploid cancer cells in later disease stages, a direct link with disease recurrence has not been firmly established. We also found that CTC-IGC have copy number profiles identical to typical CTCs and are predominantly present in the bone marrow rather than in peripheral blood.

Through single cell copy number profiling and the isolation of progeny from individual polyploid cells, we demonstrate that the polyploid cancer cell phenomenon represents a change in cell state.

Single-cell copy number profiling shows that the copy number ratios in patient CTC-IGC as well as chemotherapy induced polyploid MDA-MB-231 and PC3 cells that survive treatment are identical to those in their paired non-polyploid samples. This indicates that these cells, either identified as patient CTCs or those that survive in the days following therapy release *in vitro*, undergo multiple rounds of WGD without any additional copy number alterations. These findings provide crucial insights into the dynamics and genetic stability of the polyploid cancer cell state.

Obtaining proliferative progeny proved challenging; after three months of culturing single isolated polyploid cells, we successfully derived only one proliferative progeny clone (1/1,440). This outcome is significant for two main reasons: first, it demonstrates that polyploid cancer cells can give rise to progeny, but second, the extremely low success rate underscores why these cells have historically been understudied. To enhance our understanding, future research should employ high-throughput techniques to isolate larger numbers of single cells, such as tens of thousands, which may prove critical in understanding the roles of non-proliferative polyploid cancer cells and assessing their capabilities at reinitiating cell division to give rise to progeny.

Additionally, slight variations in the copy number profiles, such as a 3p gain observed in the progeny-1 clone, hint at genomic evolution. Further studies should explore this genomic evolution in different progeny clones once they are sufficiently collected to understand the dynamics of genomic re-organization in these cells.

Through *in vitro* single cell transcriptomics, we further provide evidence that polyploid cancer cells display a convergent phenotype between MDA-MB-231 (breast cancer) and PC3 (prostate cancer) model systems. Despite being induced with chemotherapies with contrasting mechanisms of action (cisplatin and docetaxel), the different tumor models displayed a shared polyploid signature of upregulating 309 common genes. This convergence reveals significant insights into the biological features of polyploid cancer cells.

In our observations, approximately 50% of polyploid cancer cells remained attached to the culture flask in a non-proliferative state during single cell progeny outgrowth experiments. Polyploid cancer cells have been identified to progress through the cell cycle but do not proliferate (i.e., endocycling or cytokinesis failure occur before mitosis) [55]. This is hypothesized to be a protective state of the cells that affords protection from therapeutic stressors. This phenomenon aligns with our identification of ZNF697 and NPAS2 as two transcription factors significantly enriched in the convergent polyploid gene set that were previously identified to be upregulated in cells that were in a non-proliferative state and began re-initiating cell division [47]. This suggests that some polyploid cells profiled on day 10 post therapy release may be attempting to re-initiate proliferation since the chemotherapy has been removed. This finding is further supported by a higher percentage of cells at 10-DPT expressing more markers at the M- phase of the cell cycle (Fig. S8-9). Future research should explore the roles of ZNF697 and NPAS2 in polyploid cancer cells and their implications for disease recurrence in progeny cells.

The convergent surviving cell gene set we identified indicated that pro-survival and anti-apoptotic pathways, such as TNFa via NFkB, PI3K-AKT, and mTOR signaling, are upregulated in polyploid cancer cells [57–59]. Among the genes identified in these pathways, TNFRSF9, HOMER1, and LRP1 were identified as putative survival genes and were found to be upregulated at the RNA and protein levels [48–54, 60–64]. Notably, these protein markers were retained in the PC3 progeny-1 clone, suggesting their upregulation in cells that survive chemotherapy. Additionally, a subset of CTCs, including both polyploid and typical CTCs, tested positive for TNFRSF9, HOMER1, and LRP1 at the protein level. Of note, Patient 5, who experienced the shortest progression-free survival at 1.4 months, had the highest percentage of CTCs positive for the TNFRSF9 marker, indicating that this gene may play a significant role in cancer cell survival. Further, these markers identified a subset of cells with IGC that were negative in the epithelial channel. These cells may be CTCs that lost epithelial expression (i.e., EMT) and, in combination with the upregulation of the proposed survival markers, could be adept at surviving in the bone marrow. Further studies are needed to evaluate the roles of TNFRSF9, HOMER1, and LRP1 in chemotherapy resistance and as a biomarker to evaluate the emergence of therapeutic resistance.

Our investigation of polyploid cancer cells confirms the significant upregulation of hypoxia and cholesterol homeostasis pathways. Studies have shown that targeting these pathways in cell line models, including PC3 and MDA-MB-231, reduces the viability of progeny from polyploid cancer cells [23,26]. Further evidence comes from a study indicating that polyploid cancer cells accumulate lipid droplets in response to chemotherapy [56], underscoring the critical role of lipid balance as cells significantly increase in size. These findings suggest that these pathways are integral to the polyploid cancer cell state and represent promising targets for therapeutic intervention.

The *in vitro* environment of cell culture does not always recapitulate the *in vivo* nature of cancer cell biology. This makes it difficult to speculate how polyploid cancer cells interact with their neighboring malignant cells and the surrounding stroma.

Translating the findings of TNFRSF9, HOMER1, and LRP1 as resistance markers in an *in vivo* model is a critical next step. Future studies should employ mouse models or patient derived xenografts and stain for these biomarkers to understand their prominence *in vivo*. Further studies should also isolate polyploid cancer cells through nuclear density to further understand their cellular phenotypes in tumor tissue.

While patient results are promising, they also have limitations. This study focuses on late-stage patients with disseminated CTCs in the bone marrow and blood. The evaluated cohort comprised advanced-stage patients whose previous treatment regimens had failed. To minimize biases associated with late-stage disease and to better understand initial treatment responses and their role in inducing polyploid cancer cells, future cohorts should include patients undergoing their first rounds of therapy. One concern is that CTCs in peripheral blood are typically found in later disease stages, potentially biasing our patient population towards later stages. Obtaining samples from tissue, blood, and bone marrow could address these concerns and provide valuable insights into the role of polyploid cancer cells in dissemination, initial response to therapy, and disease evolution.

## Acknowledgements

We would like to thank the patients and their caretakers, including those on active duty and veterans, for participating in this study, without whom this research would not have been possible. We thank the clinical and research consent teams MDAnderson and the VA clinics for supporting the enrollment of patients and sample collections. Components of figure 2 was created using Biorender.com.

## Declarations

### Ethics Approval and Consent to Participate

The study was conducted according to the guidelines of the Declaration of Helsinki and approved by the Institutional Review Board of the University of Southern California Keck School of Medicine, MD Anderson (NCT01505868), and the VA of Los Angeles (PCC 2015-090980). Informed consent was obtained from all subjects involved in the study.

### Data Availability

Cell line scDNA-seq (GSE270567) and scRNA- seq (GSE270568) are available through GEO. Patient scDNA-seq is available upon reasonable request. Image data is available upon reasonable request for cell lines and patients.

### Materials Availability

If interested in using the High Definition Single Cell Assay please contact CSI-Cancer.

### Code Availability

Image analysis code is freely available at https://github.com/aminnaghdloo/slide-image-utils. Downstream analysis scripts (DNA-seq, RNA-seq, image quantification) are available upon request.

### Author Contribution

Conceptualization: M.J.S., R.K.P., K.J.P., S.R.A., and J.H.; Methodology: M.J.S., A.N., R.K.P.; Software: M.J.S., A.N.; Formal analysis: M.J.S.; Investigation: M.J.S. and M.K.; Resources: P.K. and J.H.; Writing - Original Draft: M.J.S., J.H, and S.R.A.; Writing - Review & Editing: M.J.S., S.R.A., R.K.P., M.L., M.K., A.N., K.J.P, P.K., A.A, A.Z-W, P.C.; Visualization: M.J.S.; Supervision: K.J.P., S.R.A., J.H.; Project Administration: K.J.P., S.R.A., J.H.; Funding Acquisition: P.K., K.J.P., S.R.A., J.H., M.L.; Patient Accrual: M.L, R.C, I.P.G., A.A, A.Z-W, P.C.

### Consent for Publication

All the authors agree to publish this paper.

