## Supplementary methods and figures for "Polyploid cancer cells reveal signatures of chemotherapy resistance"

**Single cell picking**: Eppendorf TransferMan NK2 micromanipulator was used to collect the cell of interest in a 20 μM micropipette (for control cells) or a 100 μM micropipette (for treated surviving cells). The cell was transferred to a PCR tube containing 0.2% TritonX-100 and RNAse Inhibitor. Single cells were stored in -80°C for downstream DNA or RNA sequencing.

**Immunofluroescent staining**: Slides were removed from the -80°C and immediately fixed with 2% paraformaldehyde for 20 minutes. Slides were then blocked with 2% BSA and then incubated overnight at 4°C with a primary antibody cocktail consisting of mouse IgG1/Ig2a anti-human cytokeratins (CK) 1, 4, 5, 6, 8, 10, 13, 18, and 19 (clones: C-11, PCK-26, CY-90, KS-1A3, M20, A53-B/A2, C2562, Sigma, St. Louis, MO, USA), mouse IgG1 anti-human CK 19 (clone: RCK108, GA61561-2, Dako, Carpinteria, CA, USA), and mouse EpCAM.

**Viability**: VivaFix cell viability dye (Bio-Rad catalog 1351115) was utilized to evaluate cell permeability. Cells were treated with chemotherapy and allowed to recover. Following recovery cells were lifted with 1x versene (ThermoFisher catalog 15040066), spun down, and resuspended in 1x PBS with VivaFix dye. Cells were then plated on Marienfeld glass slides, incubated at 37°C for 30 minutes, briefly washed in PBS, and then fixed with 2% paraformaldehyde for 20 minutes. Cells were then stained with DAPI, EPI cocktail, and CD45, and imaged via high content scanning.

**Single cell copy number profiling**: Briefly, single cell whole genome amplification was performed via the WGA4 kit (Sigma) and NEB Ultra FS II was used for library preparation with 50 ng of starting material. Cells were sequenced at a depth of 1-2 million reads on an Illumina HiSeq 4000. Sequencing reads were aligned with BWA-MEM to the hg38 reference. Count data was segmented via the R package DNACopy (version 1.70.0), and median segmented ratio values were reported.

### Supplementary Tables

**Table S1**: Bone marrow patient PFS data and treatment history.

**Table S2:** Convergent polyploid gene set. 309 genes that are in common between MDA-MB-231 and PC3 polyploid cancer cells.

**Supplementary Figures: S1-S13**

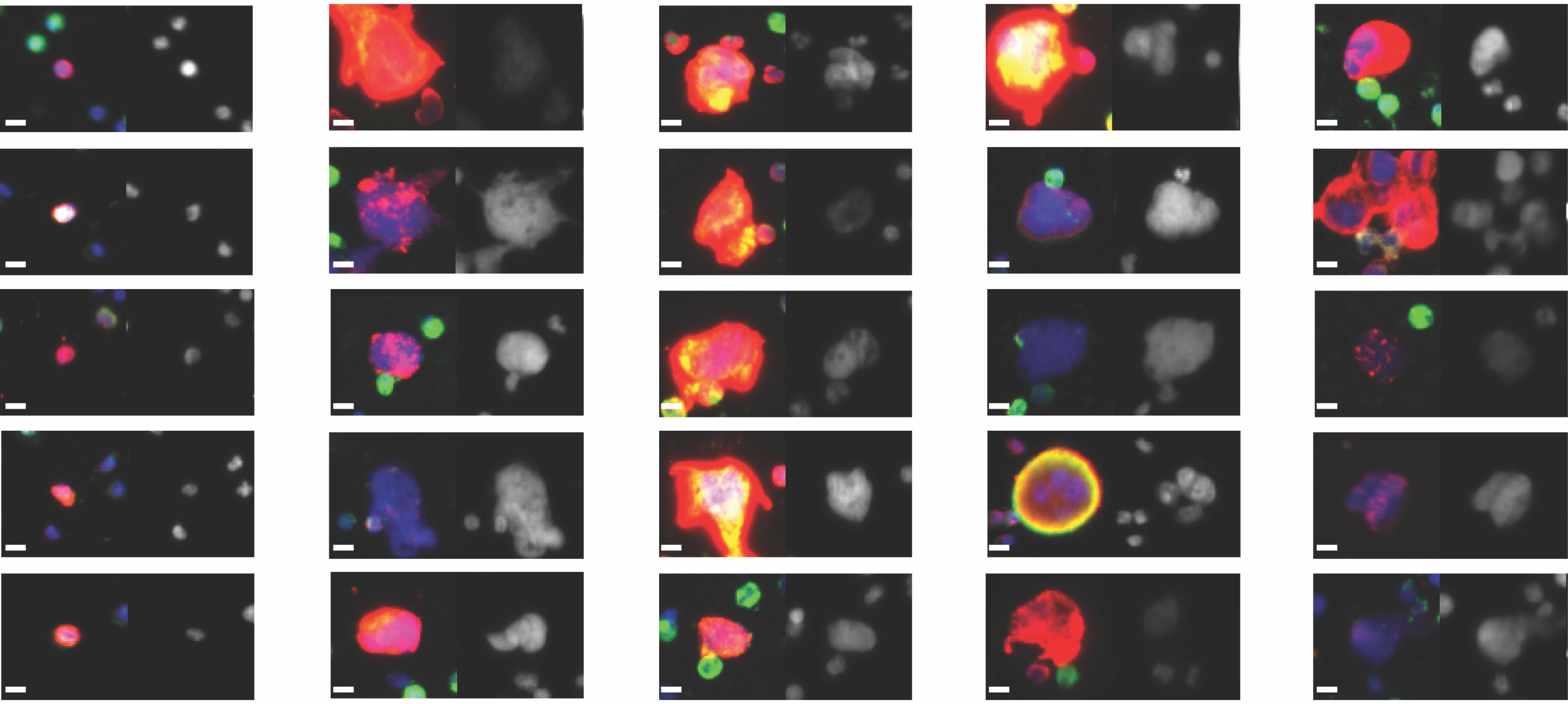

**Figure S1**: Representative gallery of typical CTCs (far left) and CTC-IGC found in bone marrow aspirate of late-stage prostate cancer patients

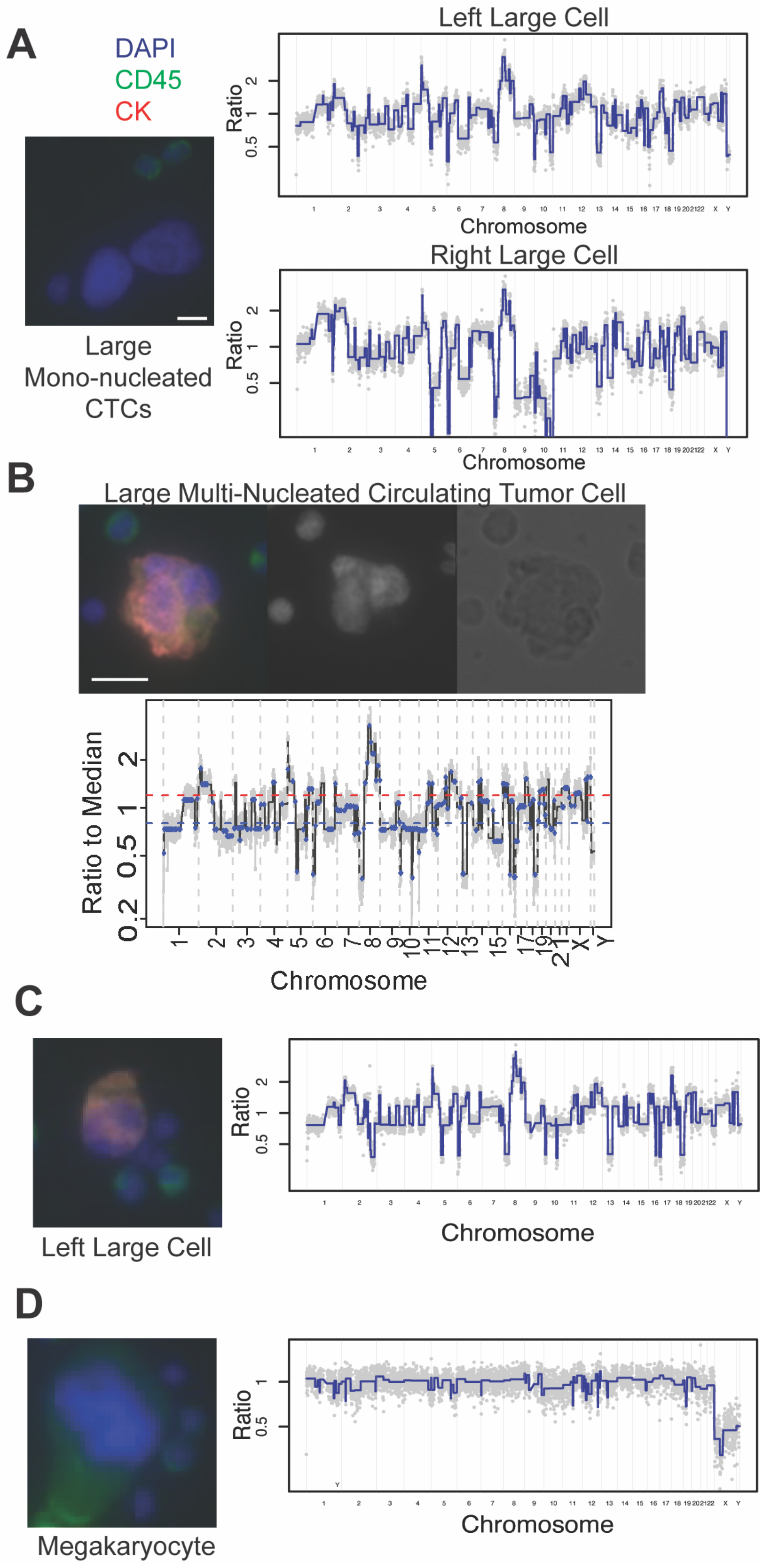


**Figure S2:** (A-C) Representative copy number profiles of CTC-IGC and (D) a non-altered megakaryocyte found in the BM of a prostate cancer patient.

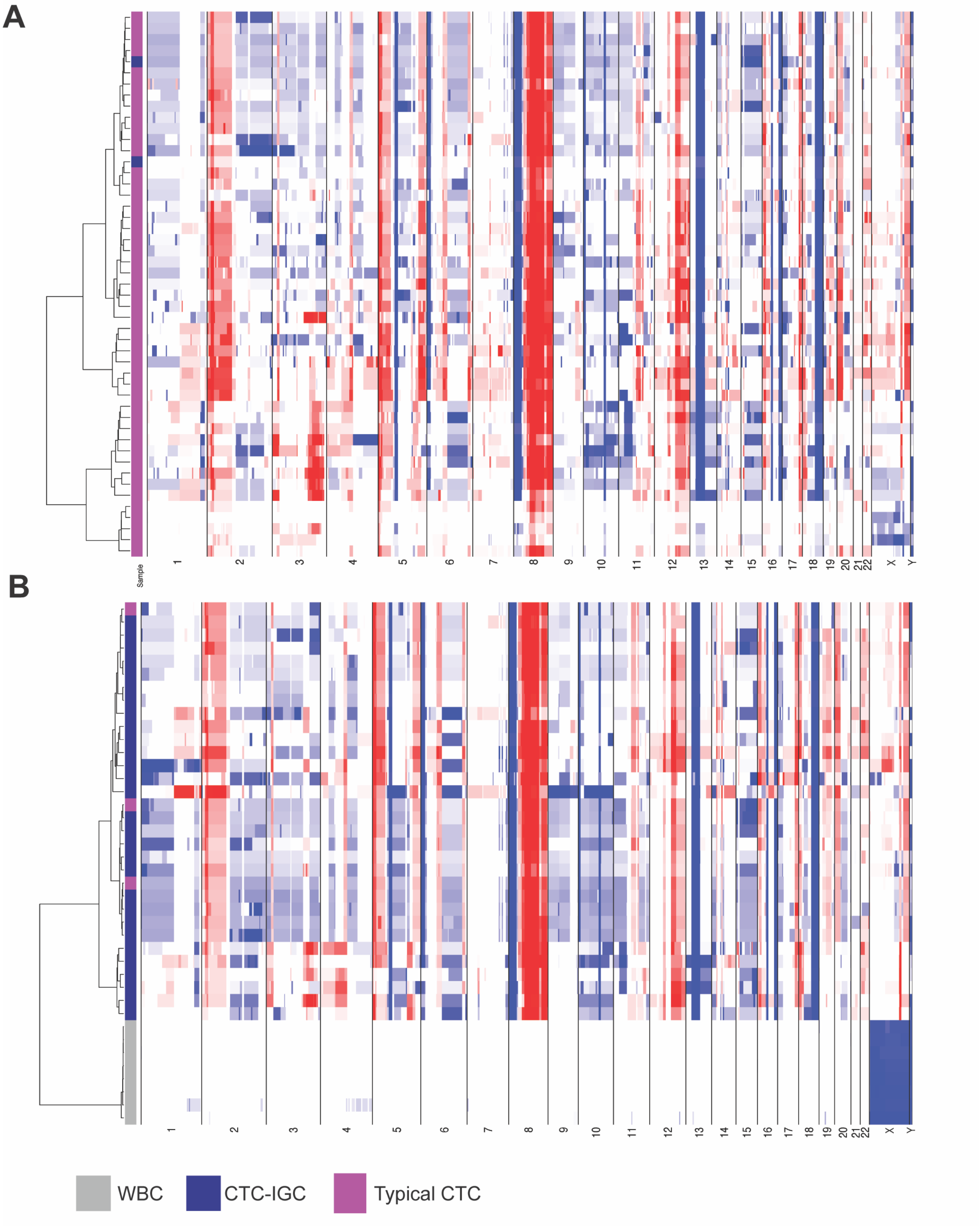


**Figure S3**: Copy number profiles from patient in figure 1 showing clonality of CTCs and CTCs-IGC. White blood cells are megakaryocytes.

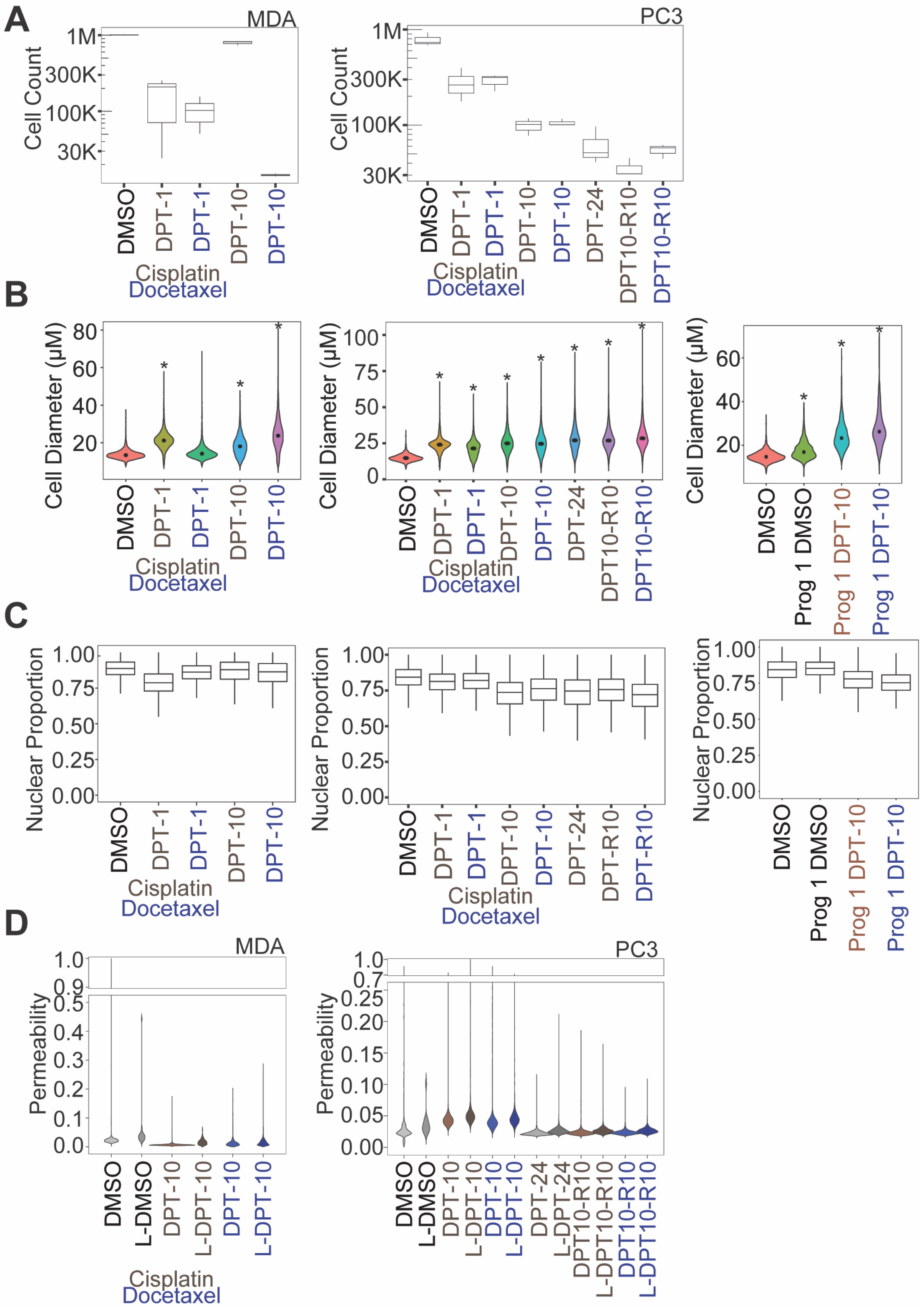


**Figure S4**: Image analysis of polyploid cancer cells. (A) Cell count for DMSO control and recovery conditions for MDA-MB-231 (left) and PC3 (right) cells. (B) Cell diameter calculations for MDA-MB-231 (left), PC3 (middle), and PC3 progeny-1 (right) cells. (C) Nuclear proportion (nuclear diameter / cellular diameter) for MDA-MB-231 (left), PC3 (middle), and PC3 progeny-1 (right) cells. (D) Permeability intensity for MDA-MB-231 (left) and PC3 (right) cells. "L" corresponds to largest 15% cells.

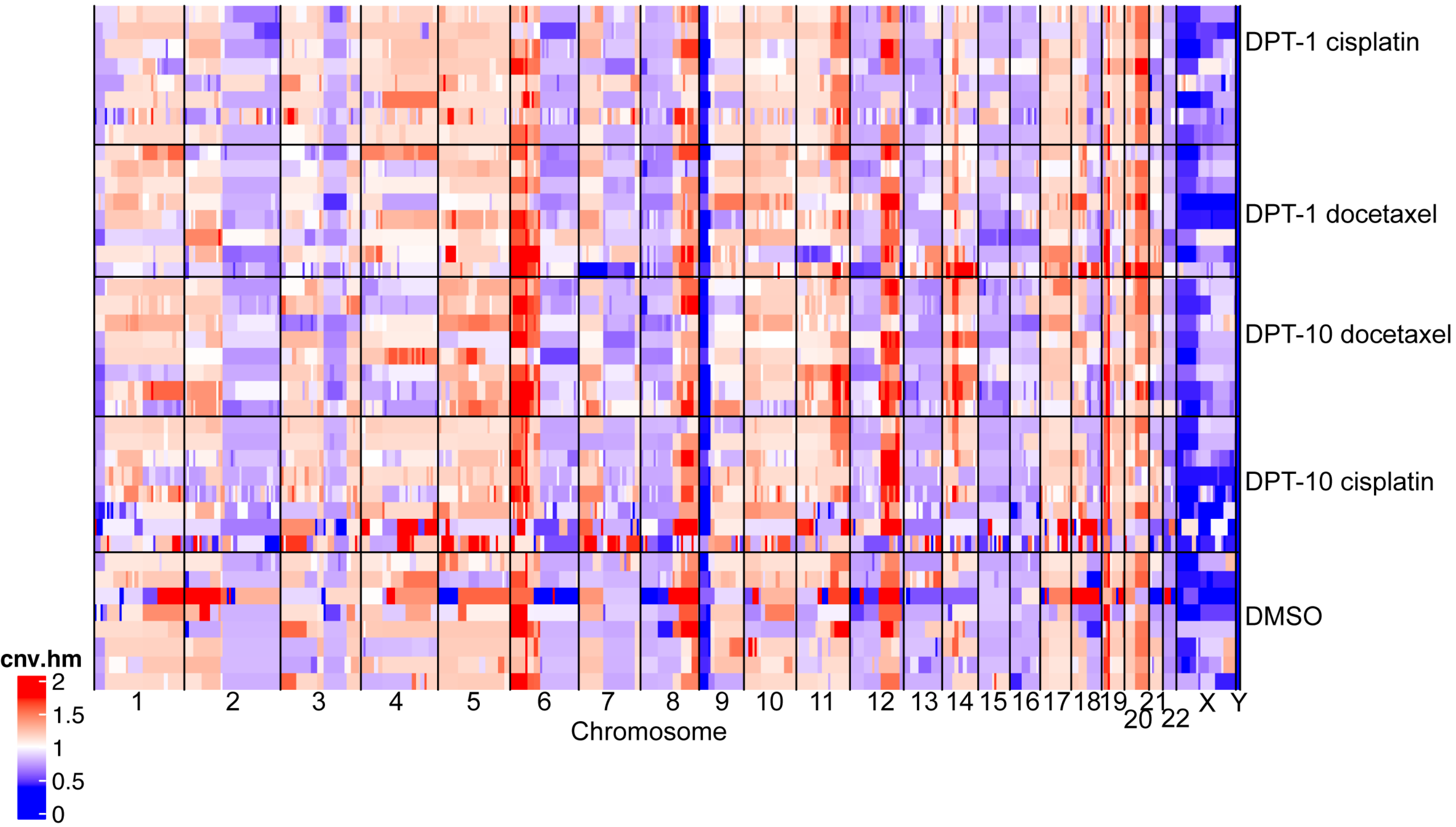


**Figure S5**: Segmented copy number ratio data for MDA-MB-231 cells.

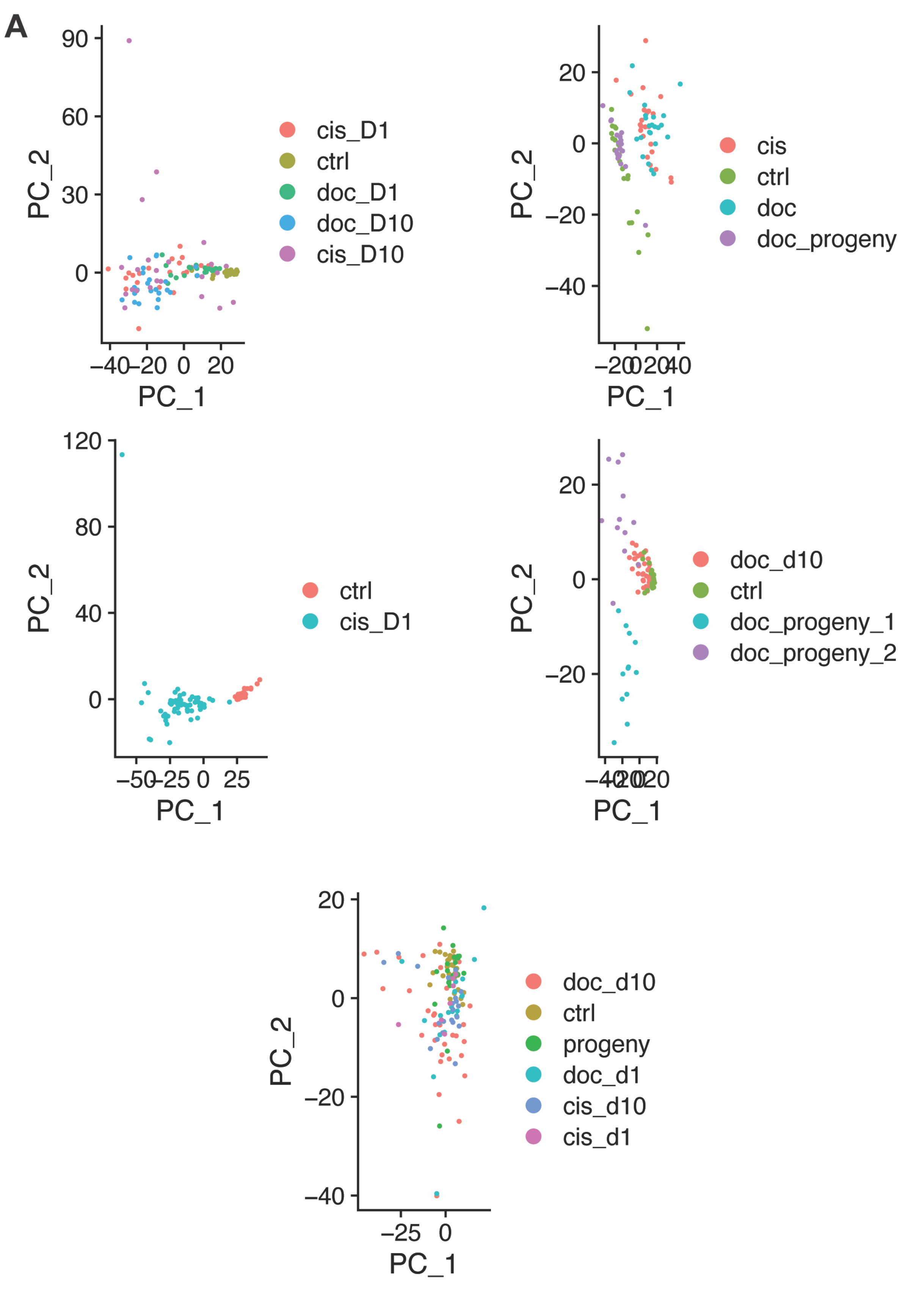


**Figure S6**: Batches of RNA from PC3 conditions.

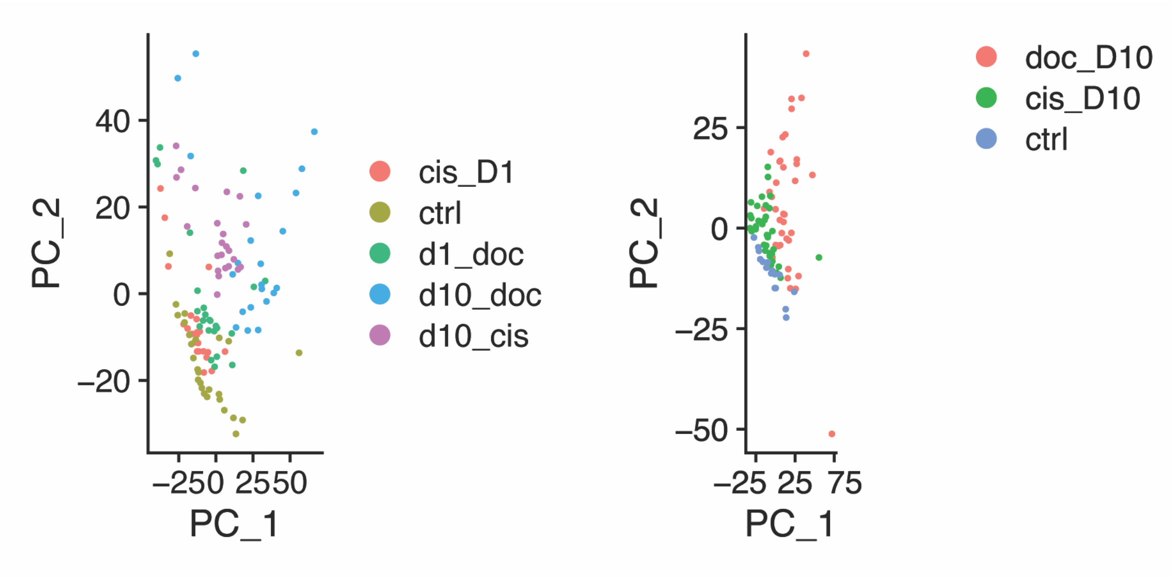


**Figure S7**: Batches of RNA from MDA-MB-231 conditions

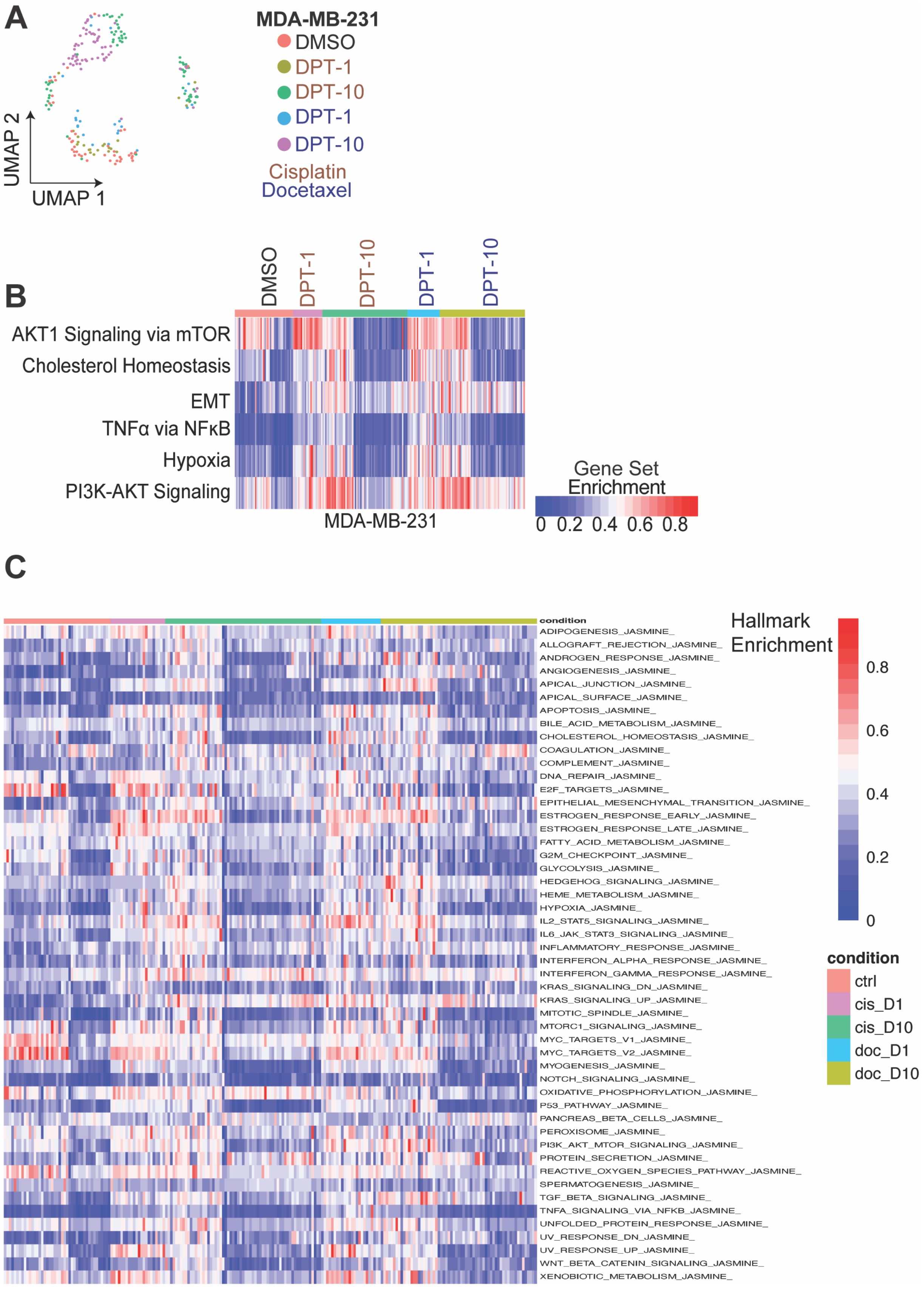


**Figure S8**: RNA sequencing of MDA-MB-231 cells: (A) UMAP visualization of MDA-MB-231 conditions. (B) Selected cell cycle hallmark pathway classifications. (C) Full single cell hallmark pathway classifications.

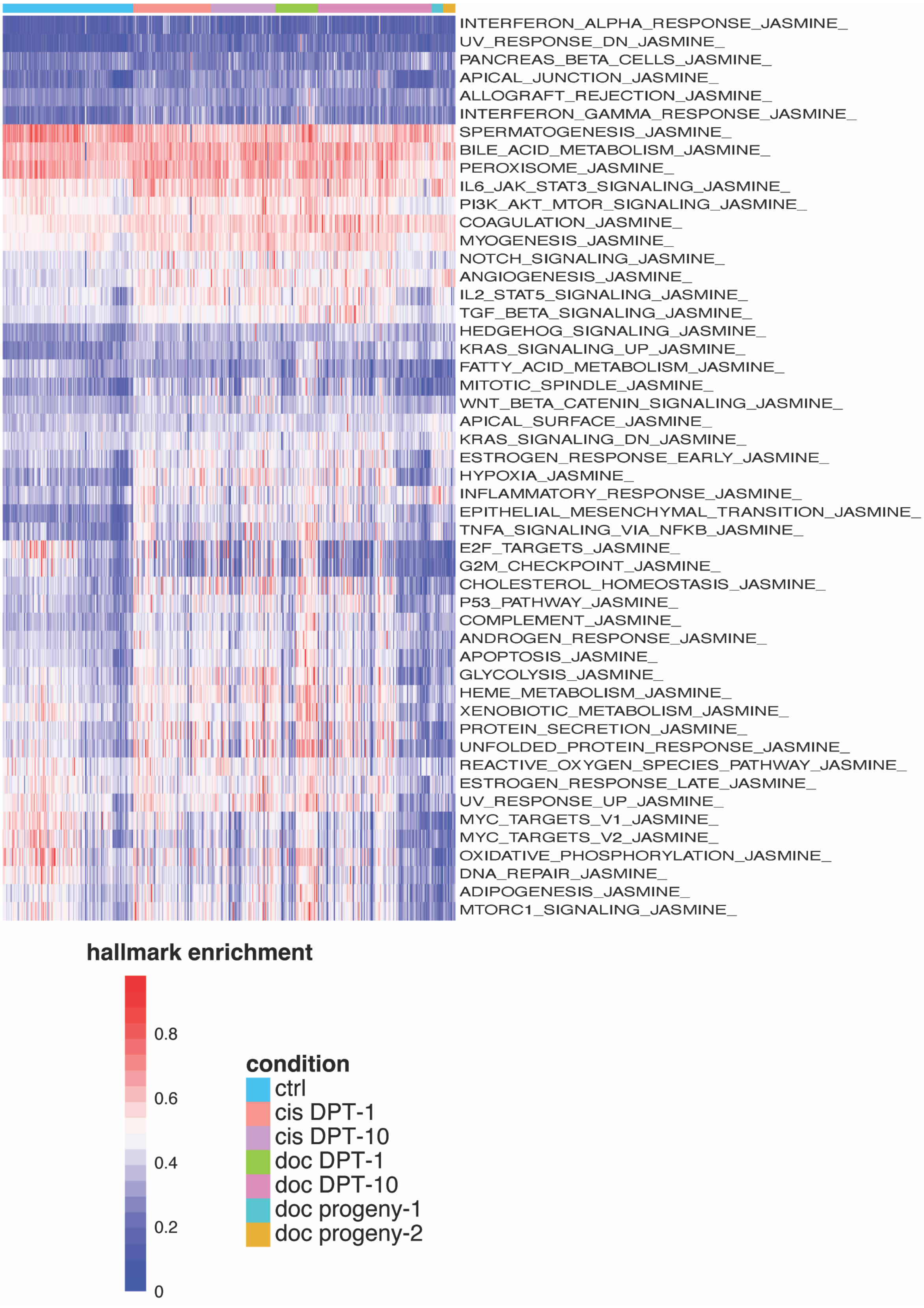


**Figure S9**: RNA sequencing of PC3 cells. Full single cell hallmark pathway classifications for each condition.

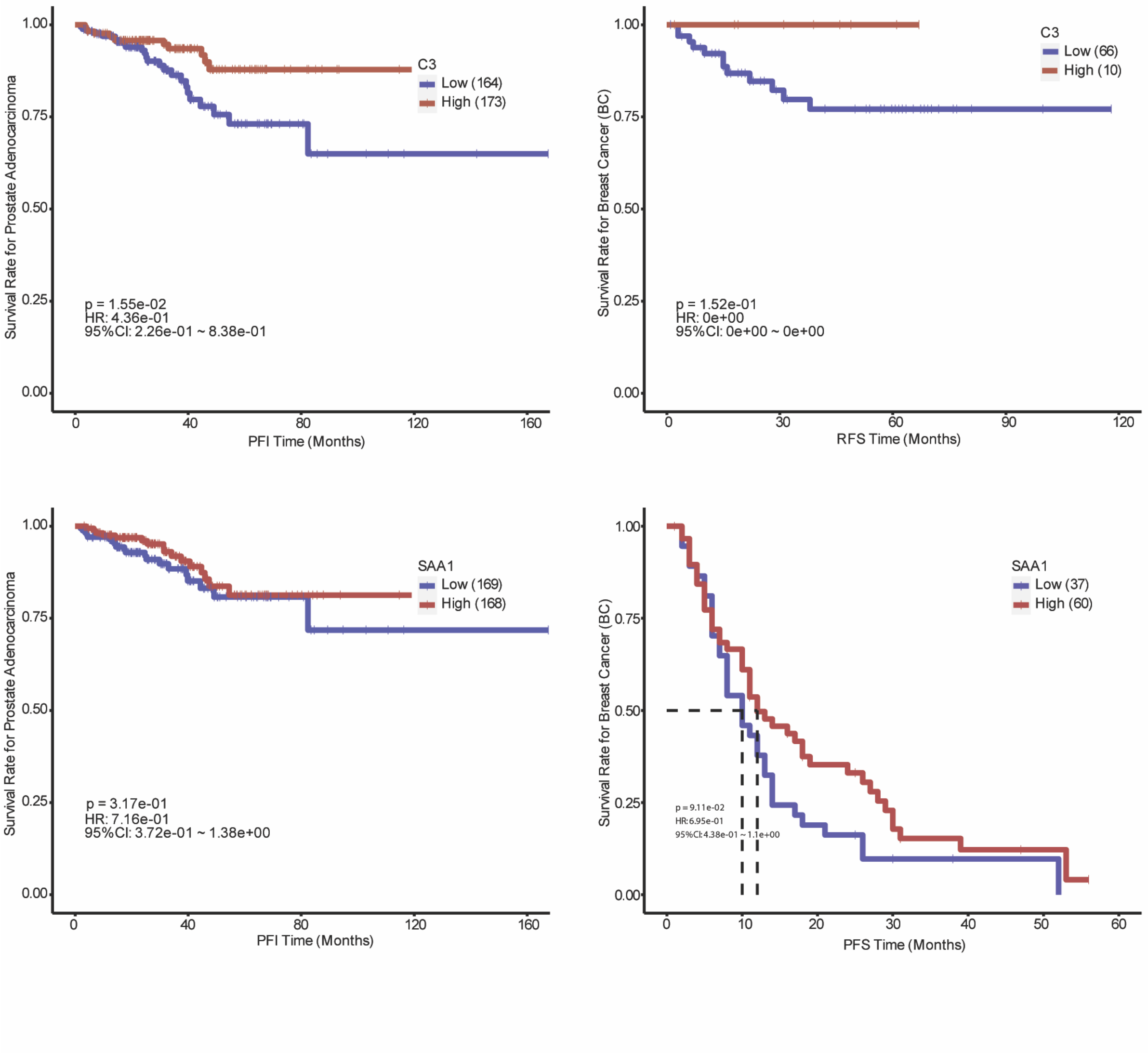


**Figure S10**: SAA1 and C3 are correlated with better PFS in breast and prostate cancer patients.

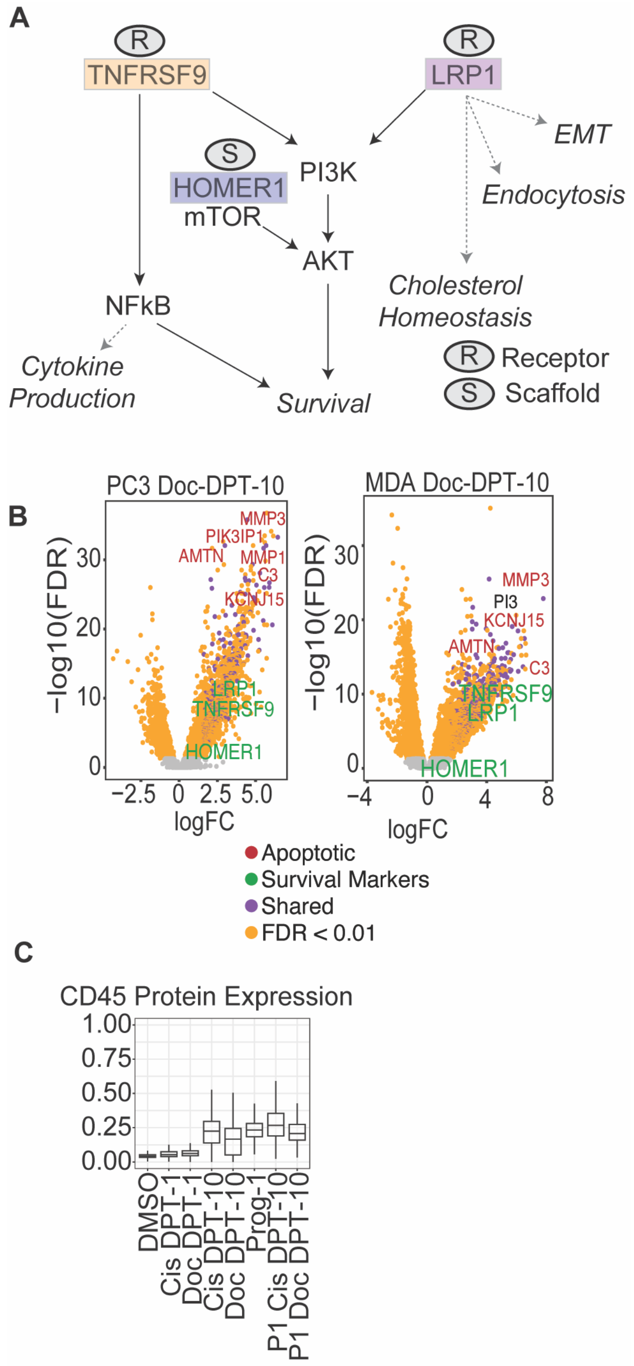


**Figure S11**: Marker supplement. (A) Hypothesized involvement of TNFRSF9, HOMER1, and LRP1 in survival pathways for cancer cells. (B) Volcano plots for PC3 Doc D10 and MDA-MB-231 Doc D10 highlight top expressed genes and marker genes. (C) PC3 CD45 protein expression increases in D10 recovered cells and is retained in progeny.

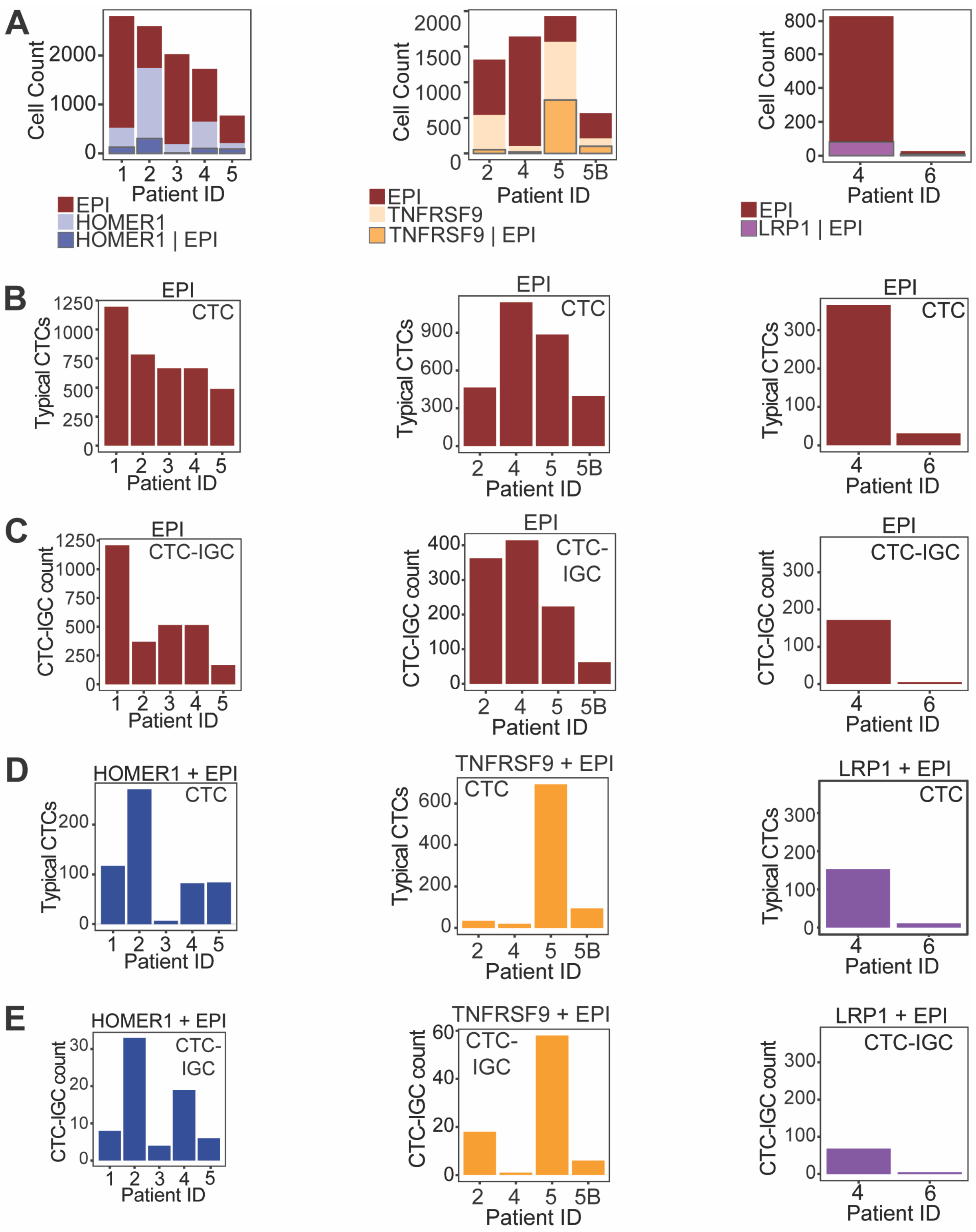


**Figure S12**: Biomarker staining in patient BM samples. (A) Total cell counts of CTCs negative for marker (EPI) and positive for marker (EPI | HOMER1, TNFRSF9, or LRP1). Cells positive for only the marker, and not EPI, cannot be confidently classified as tumor derived. (B) Signal intensity split by nuclear size. Black stars indicate gCTCs have significantly higher signal than norm-CTCs, while gray stars indicate norm-CTCs have significantly higher signal (Wilcoxon rank-sum test; $p < 0.05$).

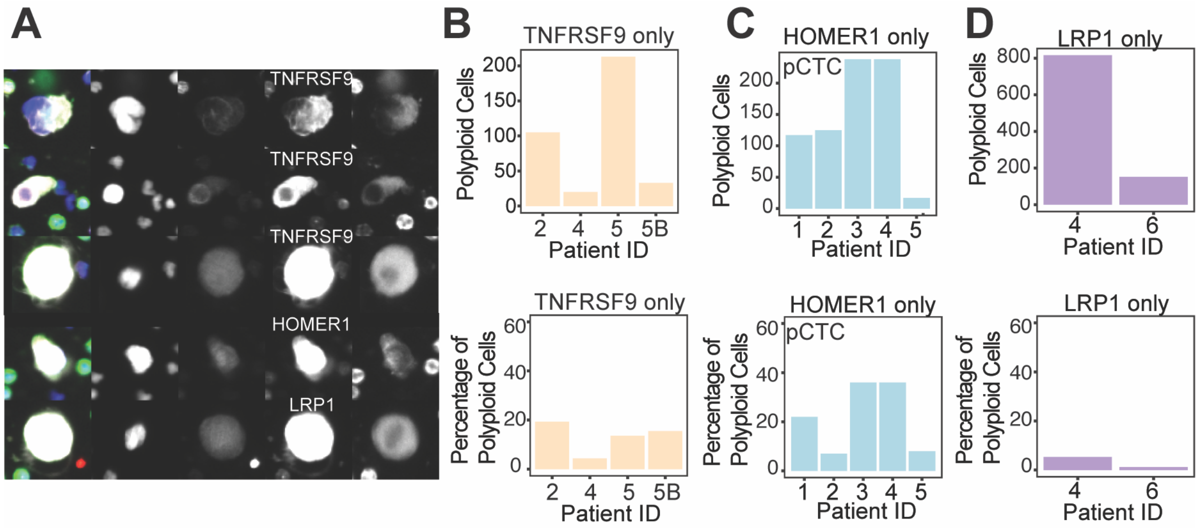


**Figure S13**: Biomarker staining in patient BM samples of polyploid cells that are positive for marker of interest but negative for EPI channel.
